# Selective enhancement of object representations through multisensory integration

**DOI:** 10.1101/740555

**Authors:** David A. Tovar, Micah M. Murray, Mark T. Wallace

## Abstract

Objects are the fundamental building blocks of how we create a representation of the external world. One major distinction amongst objects is between those that are animate versus inanimate. Many objects are specified by more than a single sense, yet the nature by which multisensory objects are represented by the brain remains poorly understood. Using representational similarity analysis of human EEG signals, we show enhanced encoding of audiovisual objects when compared to their corresponding visual and auditory objects. Surprisingly, we discovered the often-found processing advantages for animate objects was not evident in a multisensory context due to greater neural enhancement of inanimate objects—the more weakly encoded objects under unisensory conditions. Further analysis showed that the selective enhancement of inanimate audiovisual objects corresponded with an increase in shared representations across brain areas, suggesting that neural enhancement was mediated by multisensory integration. Moreover, a distance-to-bound analysis provided critical links between neural findings and behavior. Improvements in neural decoding at the individual exemplar level for audiovisual inanimate objects predicted reaction time differences between multisensory and unisensory presentations during a go/no-go animate categorization task. Interestingly, links between neural activity and behavioral measures were most prominent 100 to 200ms and 350 to 500ms after stimulus presentation, corresponding to time periods associated with sensory evidence accumulation and decision-making, respectively. Collectively, these findings provide key insights into a fundamental process the brain uses to maximize information it captures across sensory systems to perform object recognition.

**Significance Statement:** Our world is filled with an ever-changing milieu of sensory information that we are able to seamlessly transform into meaningful perceptual experience. We accomplish this feat by combining different features from our senses to construct objects. However, despite the fact that our senses do not work in isolation but rather in concert with each other, little is known about how the brain combines the senses together to form object representations. Here, we used EEG and machine learning to study how the brain processes auditory, visual, and audiovisual objects. Surprisingly, we found that non-living objects, the objects which were more difficult to process with one sense alone, benefited the most from engaging multiple senses.

## Introduction

The brain is constantly bombarded with sensory information from a host of different sources. In extracting relevant information that guides behavior, choosing which sensory information is valuable and which should be discarded is critical. This feat can be accomplished through top-down processes such as attention (Posner, 1980) or may be done more automatically by selecting stimulus features processed along each step of the feedforward processing cascade (Alais & Burr, 2004; Angelaki, Klier, & Snyder, 2009; Körding et al., 2007). In this bottom-up schema, some stimulus features may receive more weight than others given their ecological importance in guiding behavior or regularity in the environment (Laws, 2000; Patten, Mannion, & Clifford, 2017). Furthermore, an emerging literature in audition and vision suggests that these biases in perceptual weighting propagates to the level of object representations (Murray, 2006; Ritchie, Tovar, & Carlson, 2015). Given that many objects are specified not only by their unisensory features, but also uniquely identified by their multisensory signals, an open question is how multisensory cues are assembled to build object representations. Furthermore, provided that objects oftentimes share common characteristics separate from low-level features, how might potential biases in abstract object categories lead to differences in perceptual gains from multisensory integration?

One of the major categorical distinctions between objects is animacy. In vision, animate objects offer substantial processing and perceptual advantages over inanimate objects, including being: categorized faster, more consciously perceived, and found faster in search tasks (Carlson et al., 2014; Jackson & Calvillo, 2013; Lindh, Sligte, Assecondi, Shapiro, & Charest, 2019; New, Cosmides, & Tooby, 2007; Ritchie et al., 2015). Auditory studies have similarly found faster categorization times for animate objects (Vogler & Titchener, 2011; Yuval-Greenberg & Deouell, 2009). This difference may be a remnant of an evolutionary need to rapidly recognize and process living stimuli that could pose threats or be sources of sustenance (Laws, 2000). Furthermore, many inanimate objects such as cars, trains, and cellphones have not existed long enough for the brain to have developed specialized brain areas to represent them. In contrast, a number of specialized areas exist for animate subcategories, such as faces in the fusiform face area (FFA), bodies in the extrastriate body area (EBA) and voices in the temporal voice areas (TVAs) (Belin, Zatorre, Lafaille, Ahad, & Pike, 2000; De Lucia, Clarke, & Murray, 2010; Downing, Jiang, Shuman, & Kanwisher, 2001; Kanwisher, McDermott, & Chun, 1997).

To study how perceptual differences in visual and auditory categories influence their subsequent integration as audiovisual objects, it is critical to quantify neural encoding differences between objects. Representational similarity analysis (RSA) (Kriegeskorte, Mur, & Bandettini, 2008) constructs a representational space quantifying relationships between stimuli with representational distance indicating the difference in their neural signatures. A greater distance in representational space signifies more distinct neural signals between stimuli, while shorter distances signify less distinct neural signals. Studies using RSA have shown that visual and auditory objects have a clear encoding distinction between animate and inanimate categories (Cichy, Pantazis, & Oliva, 2014; Giordano, McAdams, Zatorre, Kriegeskorte, & Belin, 2013; Kriegeskorte, Mur, Ruff, & Kiani, 2008), while also showing that representational space can contract if stimuli are degraded (Grootswagers, Ritchie, Wardle, Heathcote, & Carlson, 2017) or expand in cases of increased attention (Nastase et al., 2017). Although RSA has been increasingly used to study object representations, it has not been fully leveraged to examine objects as they are often represented –as multisensory entities.

In this study, we presented subjects with auditory, visual, and semantically congruent audiovisual animate and inanimate objects while we recorded high-density EEG. Our overarching hypothesis was that greater behavioral benefits would be seen for multisensory objects and would be accompanied by an expansion in representational space as measured using RSA. A secondary hypothesis was that greater benefits would be observed for inanimate objects, given evidence that multisensory integration benefits are greatest for objects that are more difficult to process when using only one modality.

## Methods

### Participants

The experiment included 14 adults (9 men) aged 27± 4.2 years. All subjects had normal or corrected-to-normal vision and reported normal hearing. The study was conducted in accordance with the Declaration of Helsinki, and all subjects provided their informed consent to participate in the study. Each participant was compensated financially for their participation. The experimental procedures were approved by the Ethics Committee of the Vaudois University Hospital Center and University of Lausanne. EEG data for subject was removed due to poor signal quality with several artifacts present in the evoked potential response.

### Stimuli

The experiment took place in a sound-attenuated chamber (Whisper room), where subjects were seated centrally in front of a 20” computer monitor (HP LP2065) and located ~ 140 cm away from them (visual angle of objects ~ 4°). The auditory stimuli were presented over insert earphones (Etymotic model: ER4S), and the volume was adjusted to a comfortable level (~62dB). The stimuli were presented and controlled by E-Prime 2.0, and all behavioral data were recorded in conjunction with a serial response box (Psychology Software Tools, Inc.; www.pstnet.com). The auditory stimuli included 48 animate and 48 inanimate sounds from a library of 500ms-duration sounds, used in previous studies and have been evaluated in regard to their acoustics and psychoacoustics as well as brain responses as a function of semantic category (De Lucia et al., 2010; Murray, 2006; Thelen, Cappe, & Murray, 2012). The visual stimuli were semantically congruent line drawings that were taken from a standardized set (Snodgrass & Vanderwart, 1980) or obtained from an online library (dgl.microsoft.com).

### Experiment Design

Participants performed 10-13 experimental blocks (median 11 blocks) of a Go/No-Go task. Each block contained 1 audio, visual, and audiovisual presentation for each of the 96 stimuli exemplars, totaling 288 stimulus presentations per block. For half of the blocks subjects were instructed to press a button when they perceived an animate object and for the other half when they perceived an inanimate object. Animate and inanimate blocks were randomized for each subject. Auditory, visual, and synchronous audiovisual stimuli were presented for 500ms, followed by a randomized interstimulus interval (ISI) ranging from 900 to 1500ms, and participants had to respond within this 1.4-2s window. Stimuli modality was randomized for each trial (see Figure 1 for schematic). To control for motor confounds, the block instructions alternated between indicating whether the stimuli was animate or inanimate (Grootswagers, Wardle, & Carlson, 2017). Reaction times and accuracy were measured for each response. Participants did not receive feedback during the experiment.

**Figure 1.**
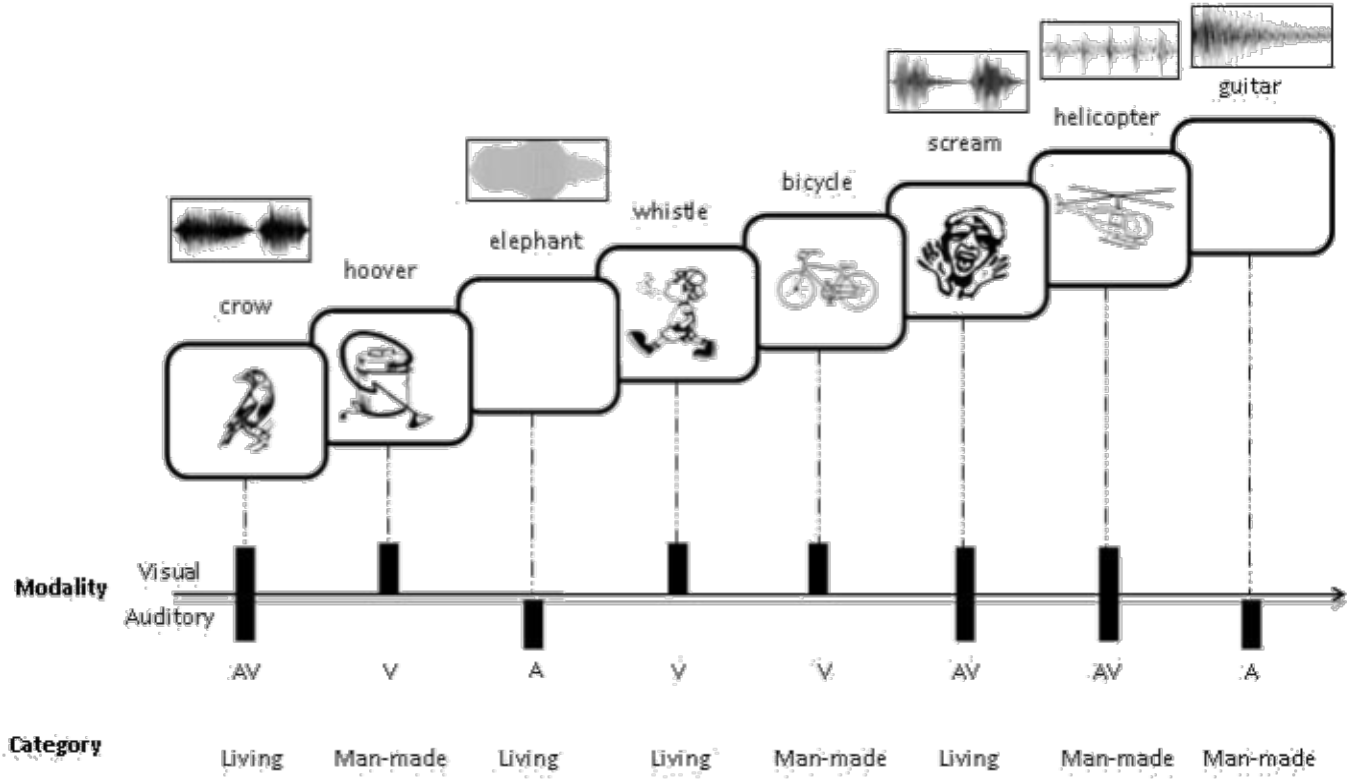
Experiment Schematic. A Go/No-Go discrimination task of animate and inanimate objects. The responses were counterbalanced such that the number of responses for animate and inanimate objects was equivalent. The stimuli consisted of 96 visual line drawings and 96 environmental sounds of common animate and inanimate objects, as well semantically congruent pairings of these objects. The sounds of animate object were non-verbal vocalizations. The stimulus duration was 500ms with a variable inter-stimulus interval of 900-1500ms.

### EEG acquisition and preprocessing

Continuous EEG was acquired from 160 scalp electrodes (sampling rate at 1024 Hz) using a Biosemi ActiveTwo system. Data preprocessing was performed offline using the Fieldtrip toolbox (Oostenveld, Fries, Maris, & Schoffelen, 2011) in MATLAB. Data were filtered using a Butterworth IIR filter with 1 Hz highpass, 60 Hz lowpass, and notch at 50Hz. All channels were rereferenced to an average reference. Epochs were created for each stimulus presentation ranging from −100ms to 600ms relative to stimulus onset. Each epoch was baseline corrected using the prestimulus period. Artifact-contaminated trials were identified and removed following visual inspection. On average 3.6± 4.7 trials were rejected. The resulting channel voltages at each time point were used for the remainder of the analysis.

### Representational Similarity Analysis

Following data preprocessing, we used CoSMoMVPA (Oosterhof, Connolly, & Haxby, 2016) and custom scripts to perform cross-validated representational similarity analysis (RSA). We used a linear discriminate classifier with 4-fold, leave one-fold out cross validation, for all exemplar pair combinations across audio, visual, and audiovisual stimuli presentations. In this procedure, trials are randomly assigned to one of four subsets of data. Three of the four subsets (75% of the data) are then pooled together to train the classifier and then decoding accuracy is tested on the remaining subset (25% of the data). This procedure is repeated a total of four times, such that each of the subsets is tested at least once. Decoding results are reported in percent correct of classifications at each time point for each exemplar pair in the time series [-100ms 600ms]. This analysis was conducted independently to build representational dissimilarity matrices (RDM) for each subject and modality over 1 millisecond increments. The RDMs were then separated into animate exemplar pairwise comparisons, inanimate exemplar pairwise comparisons, and pairwise comparisons between categories. Using these comparison groupings, mean decoding accuracies were then calculated for each modality and subject. Significant above-chance accuracies were assessed using a non-parametric Wilcoxon signed rank test, FDR (α=.025) corrected for multiple comparisons.

### Representational Connectivity Analysis

To characterize connectivity changes for different modalities and object categories, we used a combination of a searchlight analysis and representational connectivity analysis (Kriegeskorte et al., 2008). Due to this analysis being computationally-intense, data was downsampled to 100 Hz. Electrode specific RDMs, using the same procedure describe for the RSA analysis, were built by using a moving searchlight which included the electrode of interest and every immediate adjacent electrode. Electrode-specific RDMs were then correlated to each other in pairwise fashion for each electrode combination using a Spearman correlation. Note that the searchlight will change sizes depending on the chosen electrode and searchlights will overlap for electrodes and thus there will be a baseline level of connectivity in neighboring electrodes. Therefore, all connectivity measurements were compared to baseline connectivity values prior to stimulus presentation. This procedure was done for all exemplars as well as within the animate and inanimate category along the timeseries [-100ms 600ms] to compute time-resolved representational connectivity measures.

### Distance to Bound Analysis

To link neural representational space back to individual exemplar categorization times, we used a distance to bound analysis (for review see Ritchie & Carlson, 2016). Similar to RSA, this analysis represents individual exemplars as points in representational space. To decode animacy, we apply linear discriminant analysis to the representational space, defining an optimal decision boundary that separates animate and inanimate exemplars. The distance to the boundary in representational space is then computed for each exemplar across each timepoint in the timeseries [-100ms 600ms]. Next, we matched mean distance and the mean exemplar reaction time for each exemplar. We then performed a time-varying Spearman correlation between mean exemplar distance and mean exemplar reaction time for each modality.

### Model Fitting

To account for low level visual features in our visual and auditory stimuli, we constructed model RDMs and calculated their partial correlations to the neural RDM. The low-level feature auditory RDM was constructed using a Welch’s power spectral density (PSD) estimate for each of the 96 sounds. The resulting stimulus PSD was then organized into vectors and pairwise non-parametric spearman distance measurements were calculated for all exemplar pair combinations to form a model RDM. We then calculated the partial Spearman correlations between the PSD model RDM and the modality specific neural RDMs at each timepoint. An identical procedure was followed for the visual images, but instead of using PSD, image contrast was used. Note that since the images were black and white Snodgrass images, the contrast values will be equivalent to the image intensity values. In addition to these low-level feature models, we also constructed an abstract animacy category model. The animacy category model was constructed using a 0 to indicate no differences between stimuli pairs for within animacy category exemplars and a 1 to indicate complete dissimilarity for between category exemplars. This model was then also tested across modality specific neural RDMs.

## Results

### Behavior: Advantage for Animate Objects for Unisensory Presentations but not Audiovisual Presentations

Subjects were shown 48 animate and 48 inanimate auditory, visual, and audiovisual objects while they performed a go/no-go categorization task, as shown in Figure 1. We first examined behavioral differences across sensory modalities and categories, as shown in Figure 2. Figure 2A shows mean reaction times (RTs) for the go/no-go task across participants for the three sensory conditions. RTs for the auditory condition were significantly slower than for the visual and audiovisual conditions as established by the Wilcoxon signed rank test (p<0.001). Next, behavior was split by animate and inanimate categories to investigate the effects of animacy on RTs. Figure 2B shows that there are significantly (p<0.01) faster RTs for animate objects compared to inanimate objects for the auditory and visual conditions, consistent with the results from previous studies (Carlson et al., 2014; Murray, 2006; Vogler & Titchener, 2011; Yuval-Greenberg & Deouell, 2009). However, this difference is not significant for the audiovisual condition.

**Figure 2.**
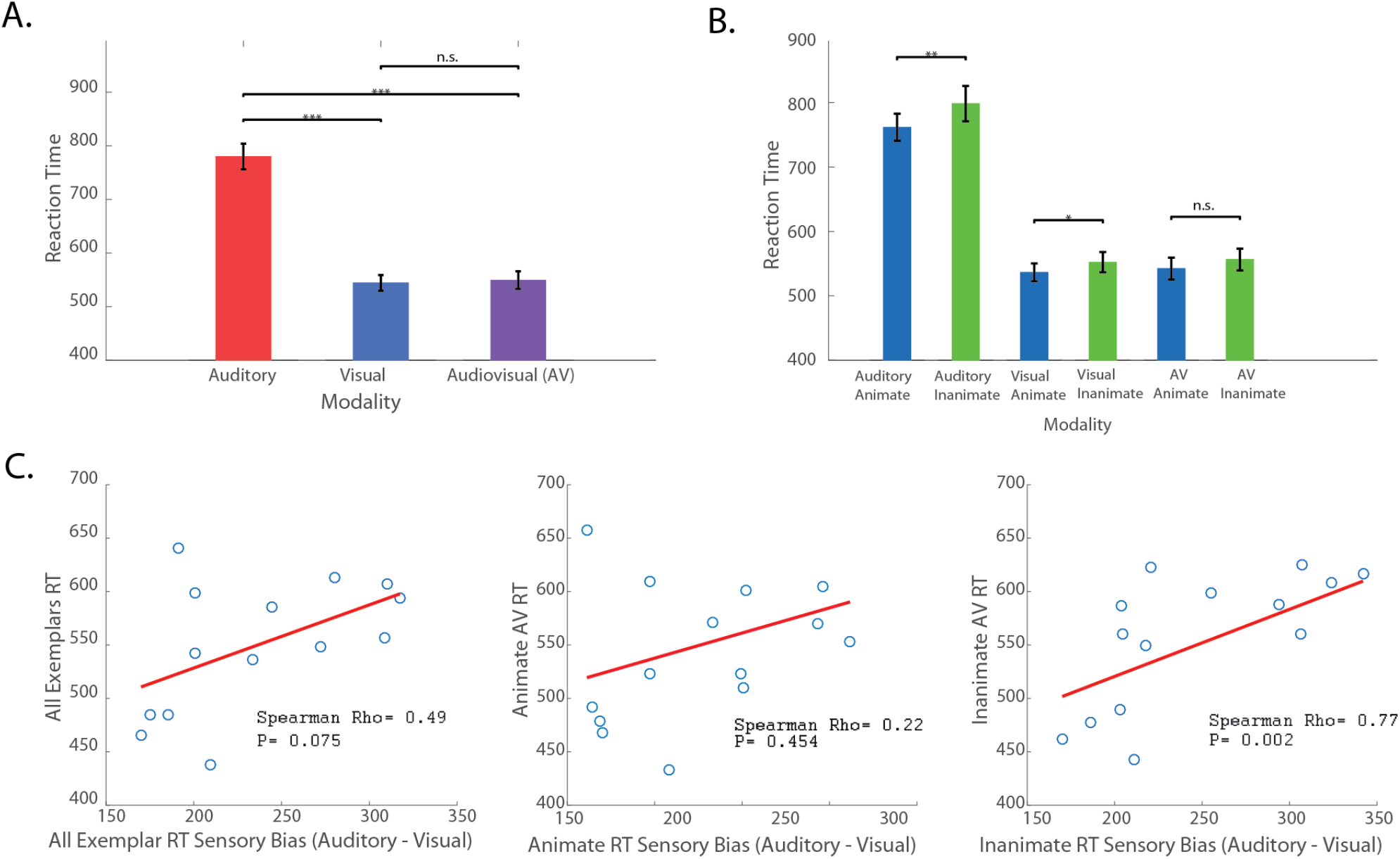
Behavior: Advantage for Animate Objects for Unisensory Presentations but not Audiovisual Presentations. (A) Reaction time (RT) results for each modality and (B) broken down by animacy. Wilcoxon signed-rank test P values (* p<0.05, ** p<0.01, *** p<0.001) above comparisons. (C) Subject sensory bias and audiovisual RT Spearman correlations across subjects for all exemplars, only animate exemplars, and only inanimate exemplars. Sensory bias is only significantly correlated to audiovisual RT for inanimate exemplars (p=0.01).

To further investigate this surprising lack of a difference in audiovisual performance, we created an index of sensory bias for each participant, operationalized as the difference in reaction times to the auditory and visual stimuli, and correlated this bias score to audiovisual RTs on a subject-by-subject basis using a Spearman correlation. Figure 2C shows that the only significant correlation between sensory bias and audiovisual RTs was for inanimate objects. The positive correlation indicates that subjects whose RTs for visual and auditory stimuli were more similar had faster multisensory RTs.

### Representational Similarity Analysis: The Influence of Sensory Modality on Between and Within Animacy Category Decoding

To investigate the neural correlates of the behavioral differences noted across conditions, we used RSA (Figure 3A-3C). Specifically, we built representational dissimilarity matrices (RDM) for each subject and modality over 1 millisecond intervals using linear discriminant analysis for each exemplar pair. From each RDM, we explored the effect of sensory modality on the distinction between animate and inanimate exemplars by calculating the mean pairwise decoding for between category pairs (e.g., dog vs. bell, dog vs. cannon). As can be seen in figure 3D, prior to stimulus onset, decoding is at chance levels (i.e., 50%), because the classifier does not have any meaningful neural data that will distinguish between category pairs. However, shortly after stimulus onset, decoding performance becomes significantly above chance (FDR corrected, p<0.025) across all three modalities. The latency of the onset of these decoding differences, defined as at least 20ms of sustained significant decoding (see Carlson, Tovar, Alink, & Kriegeskorte, 2013), was 88ms for auditory, 95ms for visual, and 60ms for audiovisual stimulus conditions. Visual and audiovisual decoding peaked at 163ms and 154ms, respectively, with higher peak decoding of 60% for audiovisual presentations compared to 58% for visual presentations. Decoding of auditory stimuli was comparatively poorer, peaking at 53% at 190ms. Note that while there were differences in significant decoding onsets, caution should be taken when comparing decoding onsets across conditions with different maximum decoding peaks (see figure 14 in Grootswagers, Wardle, & Carlson, 2017). Collectively, the results of these decoding analyses illustrate the temporal emergence of distinct neural representations for auditory, visual and audiovisual objects.

**Figure 3.**
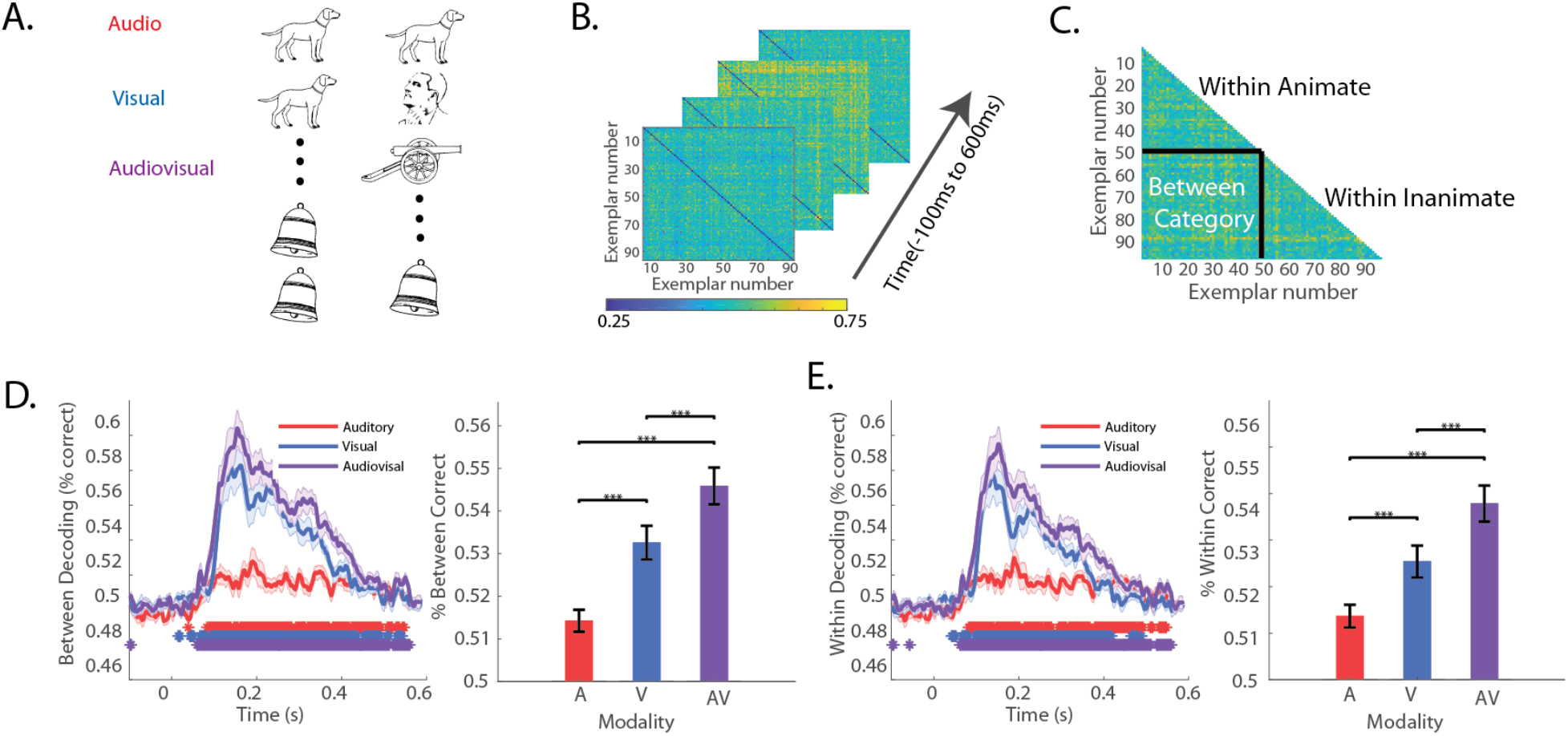
Representational Similarity Analysis: Sensory Modality Influences Between Animacy Category and Within Animacy Category Decoding. (A) RSA Schematic for pairwise decoding. Linear discriminate analysis with a 4-fold leave one-fold out cross validation was used for all exemplar pair combinations (B) Dissimilarity matrices for each of the modalities was built across time in 1 millisecond increments from pairwise exemplar classifications. (C) Mean between category and within category exemplar decoding accuracies were averaged across exemplars at each time point. (D-E) Resulting time series and summary bar plots for (D) between and (E) within categories for each of the modalities. Shaded area around lines indicates standard error across subjects. Asterisks indicate significant decoding above chance, corrected for multiple comparisons (Wilcoxon signed-rank test, FDR q = 0.025). Mean decoding across time (50ms to 600ms) for each modality with Wilcoxon signed-rank test P values (* p<0.05, ** p<0.01, *** p<0.001) above comparisons.

To statistically compare decoding performance across modalities, we computed the mean decoding for the interval spanning 50 to 600ms post-stimulus. Decoding for audiovisual stimuli was significantly higher when compared with both visual and auditory decoding (Wilcoxon signed rank test, p<0.001). Decoding for visual stimuli was higher than for auditory stimuli. These modality focused RSA results suggest that the audiovisual presentation of an object creates a more distinct representation between animate and inanimate objects when compared to either of the corresponding unisensory presentations.

We further explored whether audiovisual presentations expanded exemplar distinctions within animacy categories by calculating the mean within category pairwise decoding accuracies (Figure 3E). In this analysis, onset latencies for significant decoding for auditory, visual, and audiovisual stimuli were 89ms, 99ms, and 62ms, respectively. The corresponding peak decoding latencies were 190ms, 140ms, and 152ms. The modality-specific comparisons for within-category decoding mirrored those seen for between-category decoding, with higher audiovisual decoding when compared with visual and auditory decoding, and higher visual decoding than auditory decoding (Wilcoxon signed rank test, p<0.001). A comparison of between-category decoding and within-category decoding demonstrated higher between-category decoding for visual and audiovisual stimulus presentations (Wilcoxon signed rank test, p<0.001) but no significant difference for auditory presentations (p>.05). In sum, when compared to unisensory presentations, audiovisual stimulus presentations not only expand the representational space between animacy categories, but also make exemplars within the animacy categories easier for a classifier to distinguish.

### Category-Specific RSA: Audiovisual Presentations Selectively Enhance Inanimate Object Decoding

We further investigated representational space broken down by animacy categories to study the neural underpinnings for the observed reaction time differences between animate and inanimate categorization (Figure 4). Qualitatively, we noted that the decoding curves for animate and inanimate exemplars did not differ for auditory conditions (Figure 4A). However, this was not the case for visual exemplars, which appear to have higher decoding performance for animate exemplars when compared with inanimate exemplars. This distinction spanned the interval from approximately 100-200 ms after stimulus presentation (Figure 4B). Surprisingly, this difference is no longer apparent for audiovisual conditions. We quantified the difference between animacy categories by using the mean decoding performance during the stimulus period [50ms to 600ms]. This analysis confirmed that there was a significant animacy category difference, but only for the visual condition (p<0.05).

**Figure 4.**
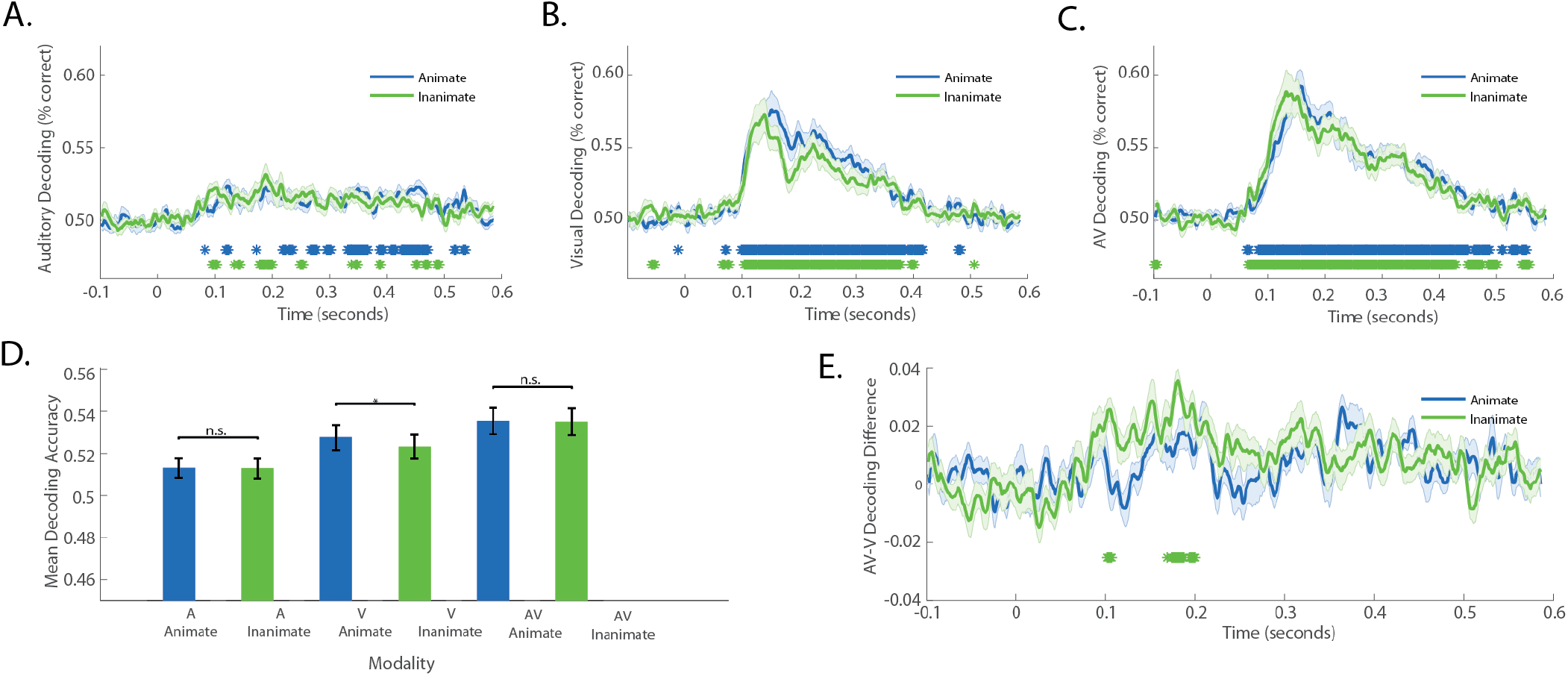
Category-Specific RSA: Audiovisual Presentations Selectively Enhance Inanimate Object Decoding. (A-C) Audio, visual and audiovisual within category decoding for animate and inanimate exemplars. Asterisks indicate significant decoding above chance, corrected for multiple comparisons (Wilcoxon signed-rank test, FDR q = 0.025). (D) Mean decoding across time (50ms to 600ms) for each modality and category with Wilcoxon signed-rank test P values (* p<0.05, ** p<0.01, *** p<0.001) above comparisons. Shaded area around lines indicates standard error across subjects. (E) The audiovisual-visual within category decoding difference for animate and inanimate exemplars with asterisks indicating significant decoding differences above 0, corrected for multiple comparisons (Wilcoxon signed-rank test, FDR q = 0.025).

Since the audiovisual condition had overall higher within category pairwise decoding than the visual condition (Figure 3E), we probed whether the lack of an animate and inanimate within-category decoding difference for audiovisual presentations was due to visual inanimate objects incurring a special benefit from audiovisual presentation. Figure 4E shows the difference between audiovisual decoding and visual decoding for animate and inanimate exemplars. Notably, the difference is significantly above zero across several timepoints between 100-200 ms post stimulus onset for inanimate objects (Wilcoxon signed rank p<0.025, FDR corrected), but not for animate objects. Furthermore, a comparison of mean decoding performance difference across the 100-200 ms time period reveals a significant difference between animate and inanimate exemplars (Wilcoxon signed rank, p=.001). Together, these results suggest that audiovisual presentations may more strongly enhance the neural representations of inanimate objects when compared with animate objects.

### Representational Connectivity Analysis: Response Patterns between Areas in the Brain are Influenced by Object Category

Given that different sensory modalities and different object classes have been shown to engage different brain networks (Braga, Hellyer, Wise, & Leech, 2017; Hillebrandt, Friston, & Blakemore, 2014), we investigated whether the pairwise decoding differences we found using RSA would also be associated with differences in mean connectivity. To carry out this analysis, we constructed electrode specific representational dissimilarity matrices (RDMs) and performed Spearman correlations across all electrode combinations to calculate a mean representational connectivity measure between electrodes. The mean representational connectivity measure is an index of how similar the representational space is between electrodes. This value is driven by two factors: spatial proximity (i.e., neighboring electrodes will have higher connectivity) and representational similarity due to stimulus features. We used the mean representational connectivity value prior to stimulus onset as our baseline, as neighboring electrodes will have shared signal even at rest (i.e. spatial autocorrelation). We found that the presentation of visual and audiovisual stimuli resulted in several timepoints that significantly diverged from baseline, and that these began at 107ms (Wilcoxon signed rank p<0.05, FDR corrected), indicating increased representational connectivity across the electrodes following stimulus presentation. In contrast, significant timepoints were not found for the auditory condition. Similar to the RSA findings, we also found that the animate and inanimate category selectively affected connectivity measurements across the different sensory modalities. For auditory objects, connectivity rose significantly above baseline for animate exemplars at 380 ms without significantly rising above baseline inanimate exemplars. Likewise, for visual objects, mean connectivity for animate objects rose above baseline at 117 with no significant timepoints for inanimate exemplars. Overall, visual animate exemplars had a greater mean representational connectivity than inanimate exemplars over the entire stimulus period (Wilcoxon Signed Rank test, p<0.001). In contrast, for audiovisual presentations, animate and inanimate categories did not demonstrate a mean difference over the stimulus period (Wilcoxon Signed Rank test, p>.05). In fact, despite a lack of an overall difference, inanimate objects diverge from baseline slightly earlier (i.e. 107 ms as opposed to 117 ms for animate objects). These results build off of the RSA analyses, and suggest that the presentation of objects in an audiovisual manner increase the representational connectivity when compared to when they are presented in a unisensory context, and furthermore that these connectivity measures increase to a greater extent for inanimate exemplars.

**Figure 5:**
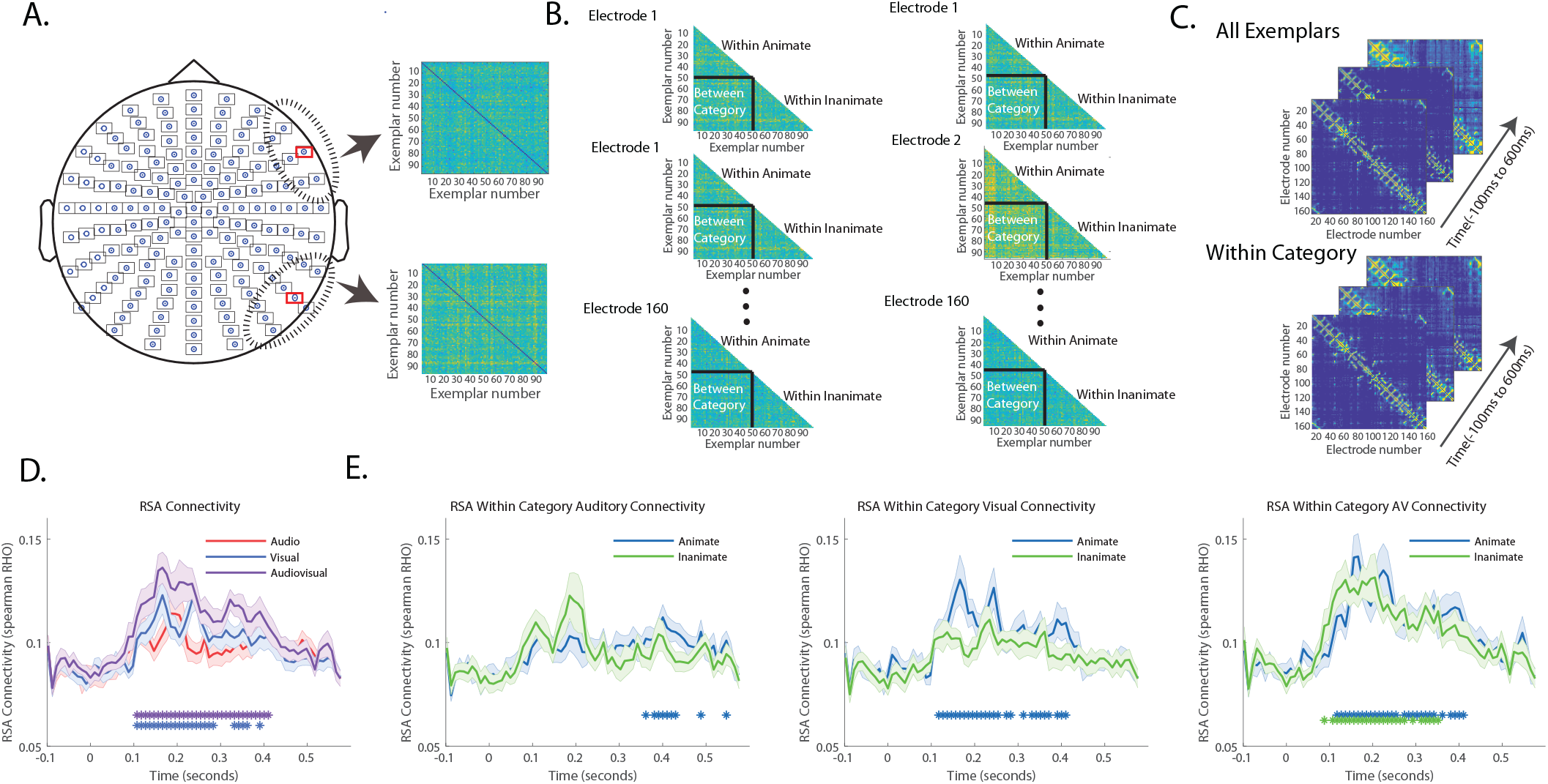
Representational Connectivity Analysis: Response Patterns between Areas in the Brain are Influenced by Object Category. (A) Moving searchlight to create electrode specific RDMs. The searchlight included the electrode of interest and every immediate surrounding electrode to produce an electrode specific RDM for each modality. (B) Each electrode was correlated in a pairwise fashion using a Spearman correlation. (C) This procedure was done for all exemplars as well as within the animate and inanimate category along the timecourse (−100 to 600ms) to build time-resolved electrode similarity matrices of representational space. The mean value of these matrices is the representational connectivity across all electrodes. (D-E) Representational connectivity was measured (D) across modalities as well as (E) within the animate and inanimate categories across modalities.

### Distance-to-Bound Analysis: Behavior can be Predicted by Exemplar Distance to the Decision Boundary in Representational Space

Having found both behavioral and neural differences between modality of presentation and animacy categories, we next considered whether the two measures were associated with one another. To do this, we computed the distance to the classifier decision boundary for all exemplars and correlated these distances with behavioral performance (i.e., reaction times). A negative correlation would denote that exemplars that are farthest away from the classifier decision boundary are those that are most rapidly categorized. Indeed, Figure 6A shows a significant negative Spearman correlation (Wilcoxon signed-rank test, FDR q = 0.025) between representational distance and reaction time at several timepoints between 100-200 ms and between 270-400 ms post-stimulus onset for both visual and audiovisual presentations. Auditory presentations did not show any significant timepoints. Figure 6B shows the corresponding scatter plot for the highest negative correlations in the 100-200 ms time window for visual and audiovisual presentations. These plots show that for both visual and audiovisual presentations, inanimate objects had slower categorization times than animate objects and were also closer to the decision boundary. Additionally, consistent with our behavioral and RSA results, inanimate exemplars appeared to show a greater shift along the reaction time and representational axes than animate exemplars when comparing between visual and audiovisual scatter plots.

**Figure 6:**
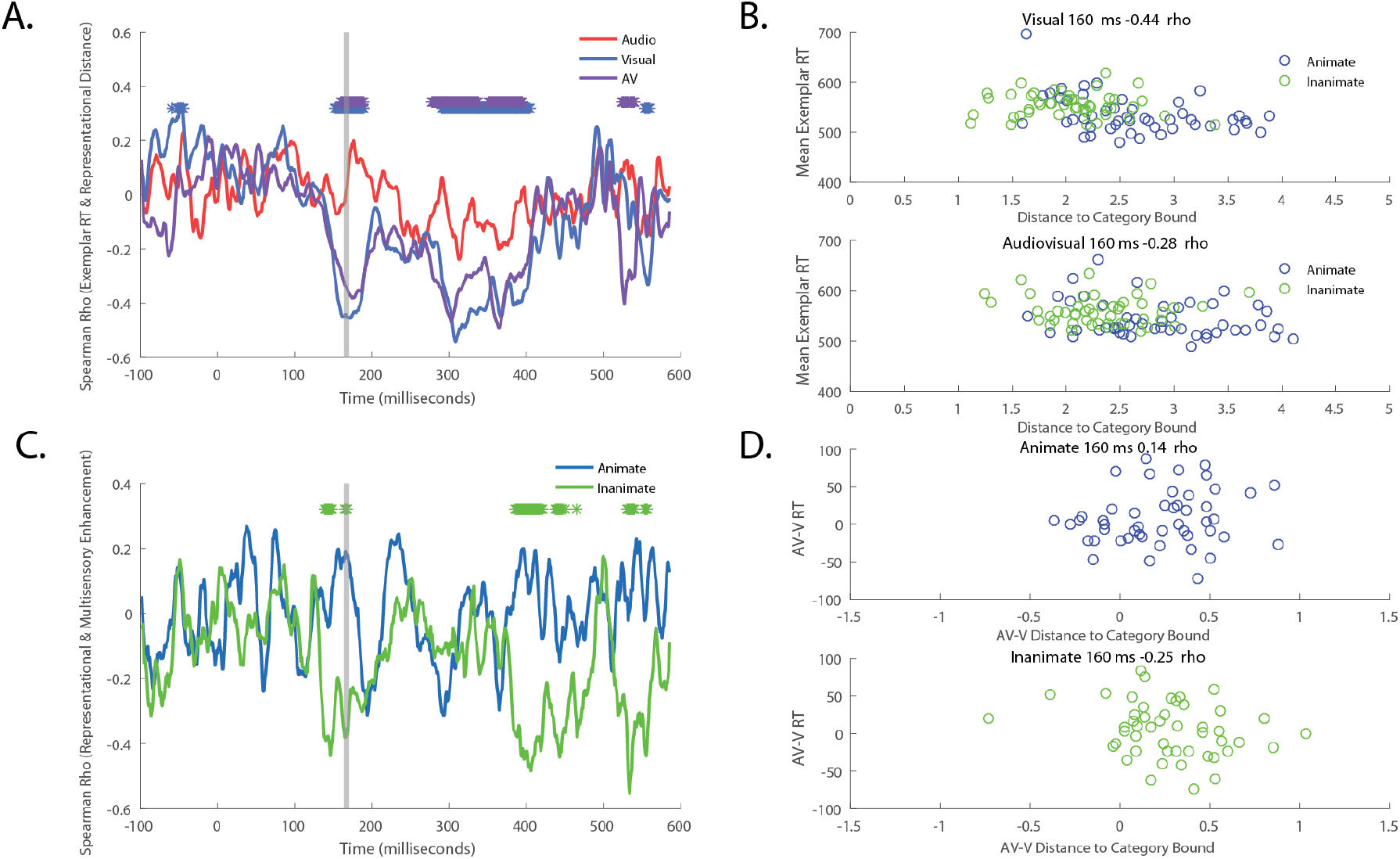
Distance-to-Bound Analysis: Behavior can be predicted by Exemplar Distance to the Decision Boundary in Representational Space. (A) Time-varying Spearman correlation between mean exemplar representational distance from animacy discriminate bound and respective average exemplar reaction time for each modality. Asterisks indicate a significant spearman correlation below 0, corrected for multiple comparisons (Wilcoxon signed-rank test, FDR q = 0.025). (B) Scatterplot for mean exemplar visual and audiovisual representational distance and RT at a significant timepoint for both modalities. (C) Time-varying Spearman correlation between mean representational enhancement (Audiovisual-Visual distance) and mean reaction time enhancement (Audiovisual-Visual RT). Note that asterisks in this plot only signify a *marginally* significant spearman correlation below 0, corrected for multiple comparisons (Wilcoxon signed-rank test, FDR q = 0.10). (D) Scatterplot for audiovisual representational and RT enhancement at a marginally significant timepoint for inanimate exemplars.

In Figure 6C, we quantified this observation by using a Spearman correlation to link the reaction time difference for audiovisual versus visual exemplars with the representational difference for animate and inanimate exemplars. A negative correlation denotes: 1) exemplars that were further away from the decisional boundary for audiovisual presentations when compared with visual presentations (positive AV-V distance value) are also the exemplars that demonstrated either more of an audiovisual RT bias (positive AV-V RT value) or less of a visual bias (negative AV-V RT value); and 2) exemplars that were further away from the decision boundary for visual presentations when compared with audiovisual presentations (negative AV-V distance value) are also the exemplars that demonstrated less of an audiovisual RT bias (positive AV-V RT value) or more of a visual bias (negative AV-V RT value). We used a less conservative FDR q of 0.10 to identify timepoints where representational distance differences show a marginally significant correlation to reaction time differences. Using this criterion, we found several significant timepoints between 100-200 ms and 370-450 ms post-stimulus for inanimate exemplars, but no significant timepoints for animate exemplars. If we calculate the mean correlation across the entire stimulus analysis epoch (50-600 ms post-stimulus) we find a significant negative correlation for inanimate exemplars (Wilcoxon signed rank, r=-0.1661 p< 0.0001) but not for animate exemplars (Wilcoxon signed rank, r=.0045 p=0.11). Figure 6D shows the corresponding scatterplot with the highest negative correlation in the 100-200 ms window for visual and audiovisual presentations at 163ms (same as figure 6B). Collectively, these results show associations between neural decoding differences and behavioral performance differences between audiovisual and visual stimulus presentations, but only when these stimuli are inanimate.

### Model Testing: Abstract Category Models Predicts Neural Activity Better than Low Level Feature Models

To account for the potential contribution of low-level features to the neural RDMs, we constructed contrast dissimilarity matrices for images and power spectral density dissimilarity matrices for sounds as shown in Figure 7. The models were correlated to each subject’s neural RDM using a Spearman correlation, and tested for significance (FDR corrected, p<0.025) along the time series ranging from 100 ms pre-stimulus to 600 ms post-stimulus. No significant timepoint emerged from this analysis, suggesting that the neural RDMs are unlikely to be a result of low-level stimulus features. In contrast, when we used an abstract model that ignored low level features and instead separated stimuli based on object animacy category, we found a significant correlation (FDR corrected, p<0.025) with the visual RDMs beginning at 118ms and audiovisual RDMs at 106ms. The animacy model was not significantly correlated to the auditory RDM, implying that the animacy distinction is not as prominent in audition.

**Figure 7.**
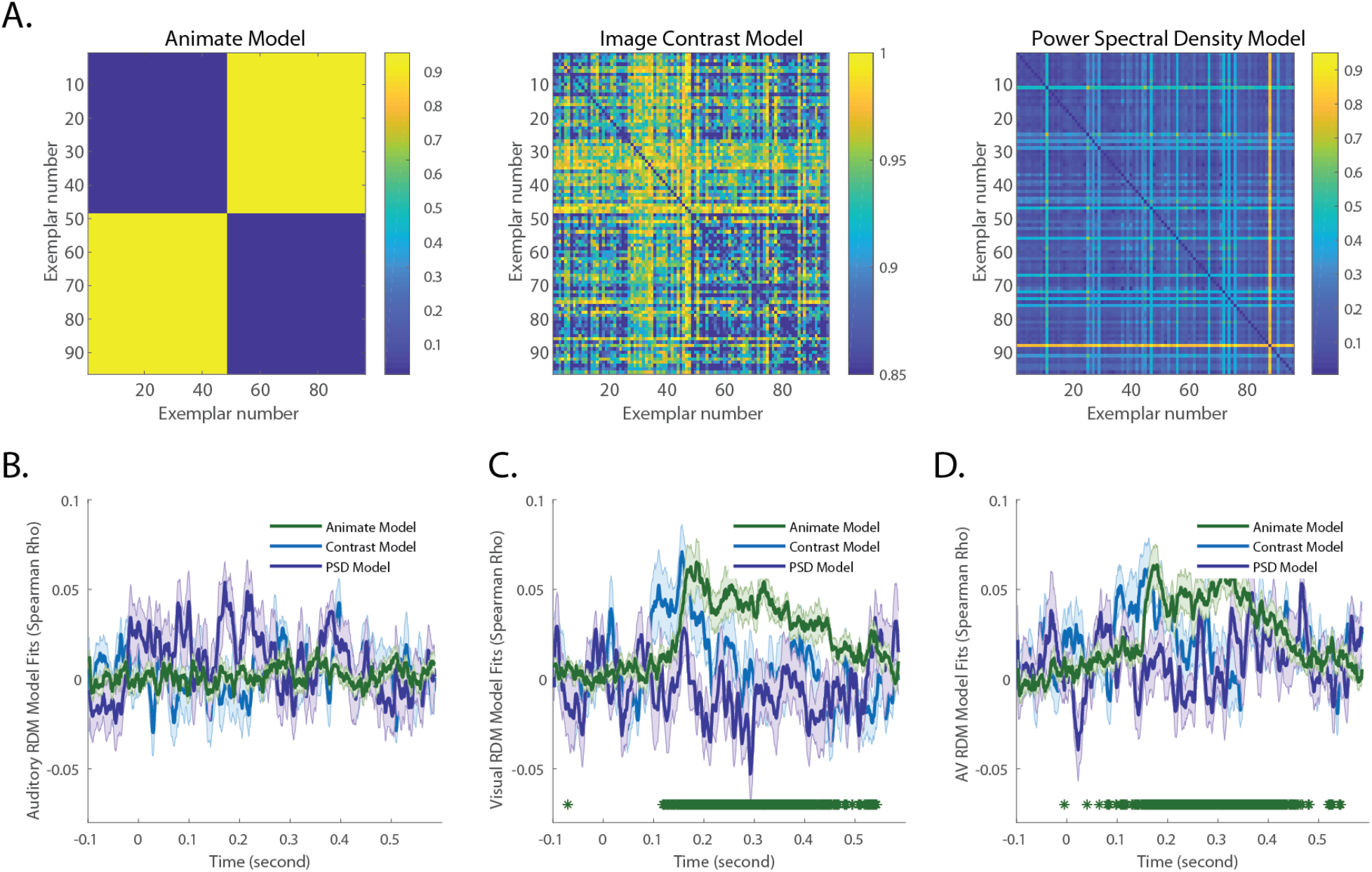
Model Testing: Abstract Category Models Predicts Neural Activity Better than Low Level Feature Models. (A) Category and low level visual and auditory feature models. The animacy category model was constructed using a “0” for within animacy category exemplars and “1” for between animacy category exemplars. For the image contrast RDM, since all images were black and white drawings, the contrast vector consisted of the intensity values of each image. The power spectral density RDM was built using a Welch’s power spectral density estimate and converted to a single vector for each sound. Each RDM was then constructed by taking the Spearman distance of each respective pairwise stimulus comparison. (B-D) Each model RDM was then tested with the (B) Auditory, (C) Visual, (D) and Audiovisual time resolved RDMs on a subject by subject basis. Shaded area around lines indicate standard error across subjects with asterisks indicating a significant spearman correlation above 0, corrected for multiple comparisons (Wilcoxon signed-rank test, FDR q = 0.025).

## Discussion

In this study, we leveraged the visual and auditory encoding bias that has been observed for animate objects over inanimate objects (Grootswagers, Ritchie, et al., 2017; Guerrero & Calvillo, 2016; Murray, 2006; Tzovara et al., 2012; Vogler & Titchener, 2011) to study how perceptual biases across object categories influences multisensory enhancement in audiovisual objects. Using behavioral measures and neural decoding, we found additional support for previous findings showing visual and auditory perceptual advantages for animate objects over inanimate objects. However, and somewhat surprisingly, we found that the advantage for animacy was not evident when objects were presented as audiovisual objects. Using RSA, we show that the lack of an animacy bias in audiovisual objects is in the context of an overall expansion of representational space when compared to visual and auditory objects. Further analysis showed that audiovisual presentations preferentially enhanced neural decoding of inanimate objects. A searchlight analysis and representational connectivity analysis showed that the presentation of inanimate objects in an audiovisual context may improve their encoding through increased representational connectivity between brain areas. We finally linked neural decoding and behavioral performance by using a distance to bound analysis and found that improved neural decoding for visual and audiovisual objects was associated with faster reaction times in the animacy categorization task. Furthermore, the decoding differences between visual and audiovisual objects was also predictive of their reaction time differences. Taken together, the results of our study provide new insights into the encoding of unisensory and multisensory objects, establishes critical links between neural activity and behavior in the context of object categorization, as well as explores potential mechanistic differences in multisensory integration for weakly and strongly encoded objects.

Although stimulus features clearly contribute to the formation of object categories, including the distinction between animate and inanimate objects, there is ample evidence that the animate-inanimate distinction transcends stimulus features and can be thought of as an abstract category distinction. The distinction is present for stimuli presented in both the visual and auditory modalities, suggesting that animacy is a general organizing principle. Furthermore, category-specific deficits in naming animate objects have been found in patients who have suffered brain damage (Capitani, Laiacona, Mahon, & Caramazza, 2003; Clarke et al., 2002; Kolinsky et al., 2002; Vignolo, 1982; Vignolo, 2004; Warrington & Mccarthy, 1987). The category distinction is preserved across species; being present in both monkey inferotemporal (IT) cortex and human IT cortex. Furthermore, the use of carefully controlled stimuli that account for stimulus features have reinforced the categorical nature of animacy (Bracci, Ritchie, & de Beeck, 2017; Ritchie & Op De Beeck, 2018). Similarly, auditory studies have also provided evidence for animacy as an abstract category distinction (De Lucia et al., 2010; Giordano et al., 2013; Murray, 2006). In the current study, we corroborate these findings by showing a significant correlation between an animacy model and neural response patterns, but a lack of any correlation between low-level stimulus features such as visual contrast and auditory power spectrum with neural response patterns.

Our study showed overall magnitude and temporal enhancement for audiovisual objects over visual and auditory objects consistent with recent findings (Brandman et al., 2019; Mercier & Cappe, 2019), and we additionally provide new insights into how audiovisual benefits selectively enhance the category of inanimate objects. Specifically, we found that the animacy bias for auditory and visual objects is absent in audiovisual objects. We hypothesized that the brain may be preferentially integrating the visual and auditory components of the more weakly encoded inanimate objects. Thus, greater multisensory integration for inanimate objects may serve to close the perceptual gap between animate and inanimate objects, consistent with the concept of inverse effectiveness (Stein & Meredith, 1993; Wallace, Ramachandran, & Stein, 2004). To test whether there were behavioral differences in multisensory integration across categories, we examined our behavioral data for a characteristic found in superior colliculus neurons: more balanced unisensory responses (i.e. smaller differences between visual and auditory stimulus) yield the greatest multisensory enhancement (Miller, Pluta, Stein, & Rowland, 2015). Behaviorally, if greater multisensory integration occurs for inanimate objects, we would also expect a stronger relationship between differences in unisensory reaction times and multisensory reaction times. In agreement, we found that smaller RT differences between visual and auditory objects led to faster multisensory reaction times for inanimate, but not for animate objects. In the same vein, the neural decoding bias for animate over inanimate objects was no longer present for audiovisual presentations. When we subtracted audiovisual decoding from visual decoding, we found that decoding was only enhanced for inanimate objects, lending further evidence that audiovisual presentations selectively improved encoding of inanimate objects.

To investigate the potential mechanism by which audiovisual presentations asymmetrically enhance the decoding of inanimate objects, we utilized representational connectivity analysis across all EEG sensors. Representational connectivity analysis has been previously used in a more limited way to assess representational similarity between two brain areas (Kriegeskorte et al., 2008). In our analysis, we used a moving searchlight consisting of each electrode and its immediately surrounding neighbors to measure the different patterns of activity for each given stimulus. By doing so, we are able to use RCA as a tool to acquire a data driven measure of how similar response patterns are topographically across the brain. We predicted that animate and inanimate exemplars might demonstrate differences in connectivity measures, as previous studies have shown increased connectivity for biologically plausible motion over mechanical motion (Hillebrandt et al., 2014). Note that in this analysis, neighboring electrodes will have shared signals simply due to proximity. Therefore, the importance of these connectivity measures is the <u>*relative*</u> difference between animate and inanimate categories. We found increased representational connectivity for animate objects when presented in vision and when compared with inanimate objects. However, much like for our RSA results, these connectivity differences were no longer present for when these objects were presented in an audiovisual context. Additionally, the connectivity increase for inanimate objects occurs within the 100-200 ms time epoch we have previously noted as the time period in which audiovisual presentations showed the greatest enhancement over visual presentations. One possible explanation for these results is that there may be increased audiovisual integration for inanimate objects relative to animate objects, leading to greater spread of neural representation across brain areas. However, the current analysis cannot exclude the possibility that the increase in inanimate connectivity for audiovisual presentations may also be due a more localized spread within electrodes in close proximity.

Next, we directly linked the neural results to behavioral results at the exemplar level by using a distance to bound approach (Carlson et al., 2014; Grootswagers, Ritchie, et al., 2017; Ritchie et al., 2015). This approach is a data-driven way of determining the relationship between neural representational space and behavioral measures (i.e., reaction times). In this analysis, we found a significant relationship between visual and audiovisual decoding distances and reaction times during two distinct post-stimulus time epochs. One corresponded to peak decoding in our RSA analysis (i.e., 100-200 ms) and the other emerged approximately 150-200ms later. These intervals potentially correspond to periods of evidence accumulation in sensory areas and brain areas responsible for decision-making, respectively (Murray, Imber, Javitt, & Foxe, 2006; Tzovara et al., 2012). We next directly correlated multisensory neural decoding enhancements to reaction time improvements. Interestingly, we found that despite an overall neural enhancement for audiovisual presentations, some exemplars showed possible effects of audiovisual interference effects. In these cases, visual decoding distances were greater than audiovisual decoding distances. These effects were largely reflected in the reaction time differences between audiovisual and visual presentation, with an overall significant negative correlation between behavioral audiovisual enhancement and neural audiovisual enhancement. These results provide evidence that the added sensory information in audiovisual presentations did not just provide the classifier with more information, but in fact provide further value for the object categorization task (Grootswagers, Cichy, & Carlson, 2018). However, it does not eliminate the possibility that added neural information was also used for other aspects of the perceptual response not tapped in the current paradigm (e.g., response confidence).

In conclusion, our study introduces new insights into the brain’s representation of sensory and multisensory information as it relates to object encoding. The greater neural encoding benefits for inanimate stimuli seen under audiovisual conditions compliments prior work, where sensory information was selectively removed from object stimuli, resulting in a selective contraction of the representational space of animate objects (Grootswagers, Ritchie, et al., 2017). Collectively, these findings show that neural representational space and the encoding of objects is impacted by both semantic congruence and stimulus modality (stimulus combinations) in a dynamic fashion. Future directions of our current work include approaches to investigate the interplay between parametrically reducing neural encoding by degrading visual stimuli while simultaneously using audiovisual presentations to enhance neural encoding. Understanding the computational framework the brain uses to maximize the sensory information it captures across sensory systems has broad implications for how stimuli perturbations and sensory integration affects object encoding.

## Acknowledgements

This work was supported by a NIGMS of the National Institutes of Health (T32GM007347). MMM is supported by the Swiss National Science Foundation (n°169206), The Fondation Asile des aveulges (n°232933), and a grantor advised by Carigest SA. We are grateful to Céline Cappe for technical assistance during data acquisition and Tijl Grootswagers for helpful comments on earlier drafts of the manuscript.

## Author’s Contributions

D.A.T., M.M. and M.T.W conceptualized the study. M.M. collected the data. D.A.T. preprocessed the data. D.A.T. performed the main data analyses and created all figures. D.A.T., M.M and M.T.W. wrote the initial draft of the paper. All authors revised the paper and approved the final version of the manuscript.

